# *In vivo* deletion of a GWAS-identified *Myb* distal enhancer acts on *Myb* expression, globin switching, and clinical erythroid parameters in β-thalassemia

**DOI:** 10.1101/2025.01.17.633540

**Authors:** Virginie Deleuze, Tharshana Stephen, Mohammad Salma, Cédric Orfeo, Ruud Jorna, Alex Maas, Vilma Barroca, Marie-Laure Arcangeli, Charlotte Andrieu-Soler, Frank Grosveld, Eric Soler

**Affiliations:** IGMM, Univ Montpellier, CNRS, Montpellier, France; Université Paris Cité, Le Kremlin-Bicêtre, France; Institut de Recherche en Cancérologie de Montpellier (IRCM), INSERM U1194, Univ. Montpellier, Institut régional du Cancer de Montpellier (ICM), Montpellier, France; Erasmus MC, Department of Cell Biology, Rotterdam, The Netherlands; CEA INSERM UMR1274, Fontenay aux roses, France; INSERM U1170, Gustave Roussy, Villejuif, France; IGMM, Univ Montpellier, CNRS, INSERM, Montpellier, France; Initiatives IdEx Globule Rouge d’Excellence (InIdex GR-Ex), Université Paris Cité, France

**Keywords:** Erythropoiesis, Thalassemia, MYB, Enhancer, globin switching, mouse KO

## Abstract

Genome-wide association studies (GWAS) have identified numerous genetic variants linked to human diseases, mostly located in non-coding regions of the genome, particularly in putative enhancers. However, functional assessment of the non-coding GWAS variants has progressed at slow pace, since the functions of the vast majority of genomic enhancers have not been defined, impeding interpretation of disease-susceptibility variants. The *HBS1L-MYB* intergenic region harbors multiple SNPs associated with clinical erythroid parameters, including fetal hemoglobin levels, a feature impacting disease severity of beta-hemoglobinopathies such as sickle cell anemia and beta-thalassemia. HBS1L-MYB variants cluster in the vicinity of several *MYB* enhancers, altering MYB expression and globin switching. We and others have highlighted the conserved human *MYB* -84kb enhancer, known as the -81kb enhancer in the mouse, as likely candidate linked to these traits. We report here the generation of a *Myb* -81kb enhancer knock-out mouse model, and shed light for the first time on its impact on steady state erythropoiesis and in beta-thalassemia *in vivo*.

## Introduction

Population-scale genetic screening have provided invaluable insights into genomic regions linked to human disease. Genome-wide association studies (GWAS) have highlighted thousands of genetic variants associated with a wide range of disorders and with disease severity or susceptibility. Interestingly, the vast majority of GWAS variants are located in the non-coding fraction of the genome, and are highly enriched in open chromatin regions corresponding to putative enhancers^1,2^. Enhancers are multifaceted distal regulatory elements enriched in transcription factor binding sites (TFBSs) which play essential roles in the spatio-temporal control of gene expression^3–6^. Mammalian genes are usually under the influence of multiple enhancers that can play overlapping as well as specific roles, to establish complex patterns of gene expression. Multiple enhancers, for instance, control the morphogen Sonic Hedgehog *SHH* expression, and *SHH* enhancers seem to individually contribute to SHH expression in specific tissues thereby displaying non overlapping functions^6,7^. Similarly, various non-redundant enhancers drive *GATA2* transcription factor expression in distinct hematopoietic cell types, thereby conferring precise spatio-temporal control of GATA2 during hematopoiesis^8^. On the other hand, clusters of distal enhancers forming the so-called globin locus control region (LCR), act in combination to maximize globin genes expression in terminally differentiating erythroid cells^5,9–12^. Both the alpha and beta globin gene loci, and their associated distal enhancers, have been extensively characterized through (epi-)genetic studies and sophisticated genome engineering approaches (e.g. enhancer KO, genetic and epigenetic engineering), providing a clear *in vivo* functional role of the individual enhancers and their complex interactions in the context of the 3D genome landscape^11–17^. Although a growing number of high-throughput functional studies are currently emerging^18–21^, such a deep characterization is lacking for most gene loci, leaving the *in vivo* roles of the vast majority of individual enhancers unclear.

In humans, the fetal and adult hemoglobins (HbF and HbA, respectively) are expressed in a temporally ordered fashion during erythroid development^22^. The fetal-to-adult switch in hemoglobin production is tightly linked to the developmental silencing of fetal globin genes (HBG1/2) and activation of adult type β-globin genes (*HBB*) which occurs in adult erythroid cells by two mechanisms: (i) direct binding of multiple transcriptional repressors such as BCL11A^23^, ZBTB7A^24^, NFIA/NFIX^25,26^ at the HBG1/2 promoters and (ii) distal LCR enhancers looping to the *HBB* gene promoter, mediated by the GATA1/LDB1 transcription factor complex, directly and strongly activating β-globin expression^14,16,27,28^. In some individuals, this developmental switch is only partially effective, leading to persistence of HbF in adulthood, a benign condition known as hereditary persistence of fetal hemoglobin (HPFH)^29,30^. This developmental switch has therefore attracted a tremendous amount of attention since reactivation of HbF in HPFH is known to ameliorate the symptoms of β-thalassemia (BT) and sickle cell disease (SCD), and to mitigate disease complications^31^. GWAS hits linked to HbF levels have repeatedly been mapped to three genomic loci, the *HBG* promoter itself, the *BCL11A* intronic erythroid enhancer, and the chromosome 6q23 *HBS1L-MYB* intergenic region^32–40^. We and others have shown that the SNPs occurring at the *HBS1L-MYB* intergenic interval, which are highly associated with elevated HbF levels and clinically important human erythroid parameters (e.g., red blood cell count [RBC], mean cell volume [MCV], and mean cell hemoglobin [MCH] content), fall within distal enhancers that control *MYB* expression^19,41,42^. The *HBS1L-MYB* interval harbors multiple *MYB* distal enhancers that are bound by the GATA1/LDB1 chromatin looping complex in erythroid cells. LDB1 mediates long-range enhancer interactions with the *MYB* gene and stimulates its expression in erythroid progenitors^5,41,42^. We have shown that some of the 6q23 intergenic variants decrease enhancer activity through the attenuation of the GATA1 and LDB1 TF binding and enhancer-promoter looping, resulting in reduced *MYB* expression levels^42^. The highest scoring SNPs pointed at a specific enhancer located 84kb upstream of *MYB* (*MYB* -84kb enh), which is conserved in the mouse (*Myb* -81kb enh), suggesting that this enhancer could play a specific role in HPFH and could directly be involved in elevated HbF levels. However, the several *HBS1L-MYB* 6q23 intergenic variants which are clustered in a ∼25kb region, are in linkage disequilibrium (LD), and cover an area overlapping with at least 6 putative enhancers^19,33,42,43^. Despite recent efforts to refine this regulatory region, fine-map the causative variants and narrow-down the region(s) involved^39^, this leaves the open questions of the role played by the individual *MYB* distal enhancers in *MYB* expression regulation and whether a single enhancer is causally involved in the HPFH phenotype and alleviates BT severity. To this aim, we generated a distal enhancer knock-out mouse model by deleting the *Myb* -81kb enhancer (i.e. the equivalent of the human *MYB* -84kb enhancer), and analyzed how it impinges on *Myb* expression *in vivo*, erythroid parameters, and globin switching. We further crossed the *Myb* -81kb enh KO animals with a mouse model of β-thalassemia (*Hbb*^th1/th1^)^44^ and surveyed the impact of -81kb enhancer deletion on the thalassemic phenotype.

## Results

### A unique transcription factor content demarcates the -81kb *Myb* distal enhancer

The mouse *Myb* gene is controlled by a cluster of 5 distal enhancers located from -36kb to -109kb upstream of its transcriptional start site (TSS)^41,45^ (Figure 1A). All enhancers display the typical chromatin signature of active enhancers and are enriched in H3K4me1 and H3K27ac. Examination of TFBS within the locus highlighted strong enrichment of the GATA1/LDB1 erythroid TF complex at all enhancers, as previously reported in mouse erythroid cells^27,41,45,46^. The human *MYB* locus displays a similar organization, with a cluster of distal regulatory elements located between -63 to -107kb upstream of the TSS^42^. These elements display strong chromatin accessibility (shown by ATAC-Seq signals)^47,48^, an active enhancer mark (H3K27ac) and GATA1/LDB1 binding (Figure 1B).

**Figure 1:**
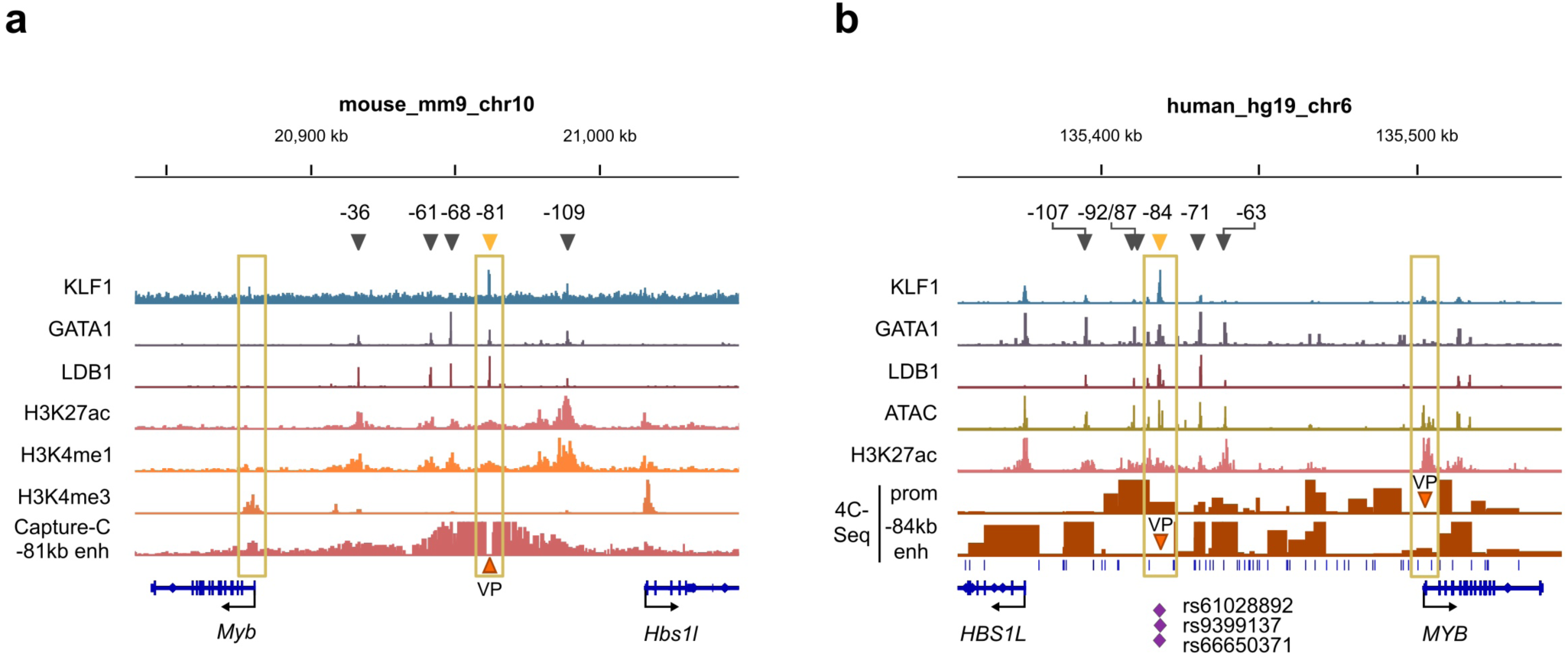
Epigenetic landscape of the MYB locus in mouse and humans. The mouse (**a**) and human (**b**) *MYB* loci are shown with ChIP-Seq tracks of select TFs from published studies in mouse (KLF1^80^, LDB1^27^, GATA1, H3K27ac, H3K4me1, H3K4me3^12^) or human cells (KLF1^21^, GATA1 and H3K27ac^19^, LDB1^42^). Human ATAC-Seq data are from^47^. Capture-C^81^ (mouse) or 4C-Seq^82–84^ data showing long-range interactions throughout the locus from the mouse *Myb* -81kb enh, and human *MYB* -84kb enh and *MYB* promoter viewpoints (VP) were performed in MEL cells (mouse) or UT7 cells (human). The BglII cut sites are shown underneath the 4C-Seq tracks (human locus). Position of the distal enhancers is indicated above the ChIP-Seq tracks (distance in kb relative to the transcription start site). The top scoring SNPs associatied with HbF levels and clinical erythroid parameters are also shown (human track).

The mouse -81kb enh sequence is orthologous to the human -84kb enh, which harbors the genetic variants most highly associated with HPFH, fetal globin levels and variations in clinical erythroid parameters^42^. Remarkably, both the mouse -81kb enh and human -84kb enh stand out as being uniquely bound by KLF1, in addition to the GATA1/LDB1 complex, rendering them unique among the locus (Figure 1A-B). Chromatin looping experiments confirm that both enhancers interact with the *Myb* promoter and first intron regions (Figure 1A-B), supporting a role in *Myb* regulation. Since KLF1 is a major regulator of erythropoiesis, this suggests that the -81kb/-84kb enh may play a specific role within the locus.

### The mouse -81kb enh/human -84kb enh are erythroid-specific enhancers

Since MYB is expressed in most hematopoietic progenitors, including hematopoietic stem cells (HSCs), myeloid and lymphoid progenitors^49–53^, we sought to determine whether the -81/-84kb enh are broadly active throughout hematopoiesis or whether they display lineage specificity. We analyzed published chromatin accessibility datasets (ATAC-Seq) from multiple primary human and mouse hematopoietic cell types^54,55^, in addition to datasets specifically focused on erythropoiesis, including terminal differentiation^47^ (Figure 2A-B). Strikingly, chromatin opening at the -84kb enh is completely absent from HSCs, non-erythroid myeloid cell types (e.g. GMPs, monocytes) and lymphoid lineages (NK, B cells, CD4 and CD8 T cells), and is restricted to erythroid committed cells (MEPs and erythroid progenitors), with low levels in CMPs (Figure 2A). This absence of chromatin opening is likely not due to inactivity of the *MYB* locus outside erythroid-committed cells as the *MYB* gene is transcribed in most of these cell types. Besides, other putative regulatory regions such as the -92kb site appear to be broadly active in hematopoiesis, with clear signs of chromatin opening in HSCs, immature progenitors (MPP, LMPP), and both myeloid (CMP, GMP) and lymphoid committed progenitors, indicating that the locus is active in these cell types. We next looked specifically within the erythroid lineage using datasets derived from *ex vivo* differentiation of CD34+ hematopoietic stem/progenitor cells (HSPCs)^47^. Chromatin accessibility at *MYB* -84kb enh becomes apparent at the early erythroid progenitor stage (Bust forming unit erythroid, BFU-E) and gradually increases up to the pro-erythroblast/early basophilic erythroblast precursor stages, and becomes progressively lost as cells fulfill terminal differentiation (late basophilic, polychromatic and orthochromatic erythroblast stages) (Figure 2B). Similar observation was done using mouse hematopoietic cells, with -81kb enh chromatin opening being restricted to erythro-myeloid cells, MEP, erythroid and megakaryocytic progenitors (Supplementary Figure 1). These observations support the notion that the conserved mouse -81kb/human -84kb enhancer displays lineage-specific activity, with erythroid (and to a lower extent megakaryocytic) specificity.

**Figure 2:**
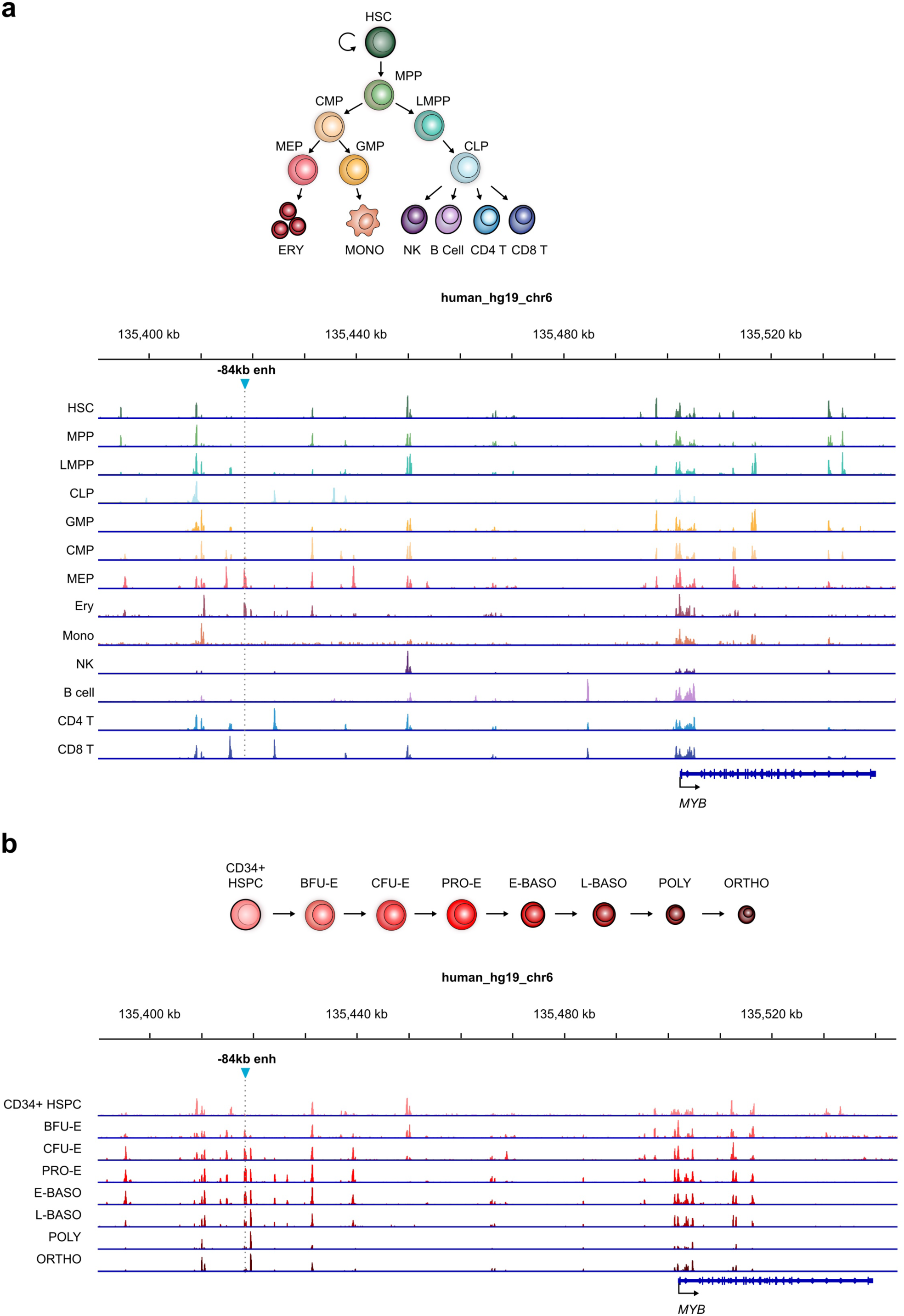
The *MYB* enhancer landscape during hematopoietic and erythroid differentiation. (**a**) ATAC-Seq signals showing open chromatin regions in the human *MYB* locus from sorted human hematopoietic populations^54^ are depicted, showing erythroid specificity of the *MYB* -84kb enhancer. (**b**) Dynamic accessibility (ATAC-Seq) and differentiation stage-specificity of the human *MYB* -84kb enhancer throughout terminal erythropoiesis (using CD34^+^-derived *ex vivo* differentiation)^47^.

### Generation of a *Myb* -81kb enh KO mouse model

The above-mentioned observations prompted us to investigate the *in vivo* role of the - 81kb enh and test whether its inactivation recapitulates the changes in erythroid parameters observed in humans harboring genetic variants at the -84kb enh region. We defined the -81kb enh as the 347bp-sized region occupied by the GATA1/LDB1 complex (Figure 3A). The corresponding genomic sequence harbors the typical GATA1::TAL1 motif i.e. CTGN(6-8)WGATAR recognized by the GATA1/LDB1 complex, and the CACCC box motifs recognized by KLF1 (Figure 3B). The -81kb enh sequence was flanked by LoxP sites by homologous recombination in embryonic stem (ES) cells (Figure 3C), which were used for blastocyst injections to generate Floxed -81kb enh knock-in mice (see methods). These mice were crossed with a mouse line ubiquitously expressing the Cre recombinase to generate *Myb* -81kb enh homozygous KO mice, referred to as -81kb enh^KO^. The -81kb enh^KO^ animals were viable and fertile, and were obtained in a Mendelian ratio (Figure 3C), indicating that removal of the - 81kb enh is not lethal.

**Figure 3:**
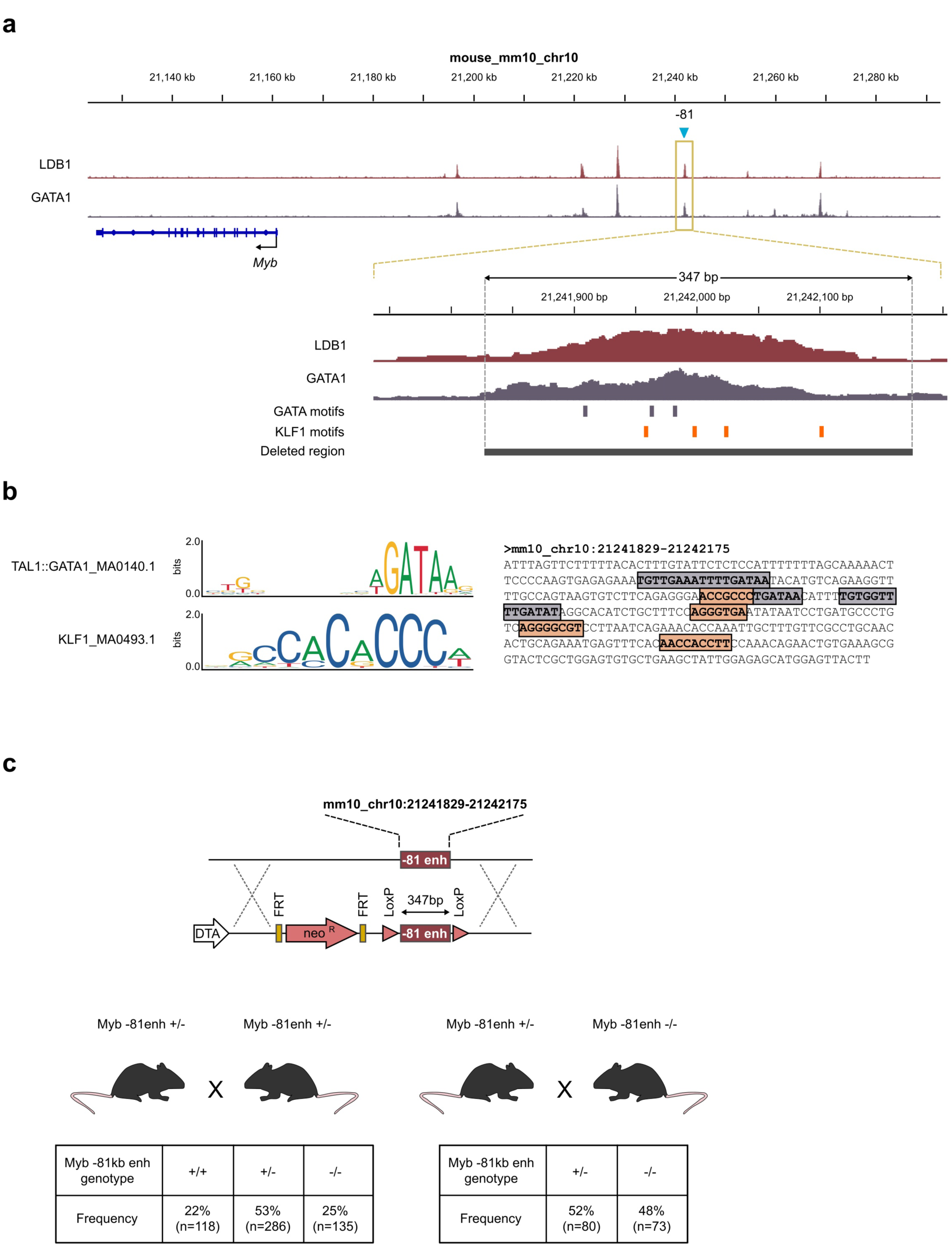
Generation of the mouse *Myb* -81kb enhancer knock-out. (**a**) The mouse Myb locus is shown with a zoom view of the -81kb enhancer, and associated LDB1/GATA1 binding site, GATA and KLF1 DNA motifs, and location of the deleted region. (**b**) Position weight matrixes of the LDB1/GATA1 complex (TAL1::GATA1_MA0140.1) and KLF1 (KLF1_MA0493.1) from JASPAR, and their position within the deleted enhancer sequence. (**c**) strategy used to knock-out the -81kb enh sequence in mouse embryonic stem cells, to generate the mouse *Myb* -81kb enh KO. The Mendelian ratios of heterozygous and homozygous animals are shown.

### Erythroid phenotype of the -81kb enh^KO^ mice

We first investigated erythroid cells development in the fetal liver (FL), representing the main hematopoietic organ at mid gestation. FACS analysis of embryonic day 13.5 (E13.5) FLs revealed decreased levels of c-Kit+ progenitors in KO animals (Figure 4A). Fine examination of differentiating erythroid populations based on the CD71/Ter119 surface markers revealed subtle but significant decrease in the most immature cells (R1, CD71^-^Ter119^-^), slightly increased R2 population (CD71^+^Ter119^-^) and a decrease in the most mature R4 cells (CD71^-^ Ter119^+^) indicating that deletion of the -81kb enh has a slight impact on erythroid maturation in FLs with a decreased or delayed output of mature red blood cell production (Figure 4A-B). We next investigated the impact of -81kb enh removal on adult hematopoietic populations. As expected from the chromatin accessibility patterns of the -81kb enh, the bone marrow immature hematopoietic compartment defined as lineage-negative (Lin^-^), Sca1-positive (Sca^+^), Kit-positive (Kit^+^) cells (LSK) containing the hematopoietic stem/progenitor cells displayed no significant alterations (Figure 4C and Supplementary Figure 2). Within the LSK cell population, both long-term, short-term HSCs (LT-HSCs and ST-HSCs, respectively), and multipotent progenitors (MPPs) were found in equal amounts in wild-type (WT) and KO animals, indicating that the stem and early progenitor cells were unaffected. Common lymphoid (CLPs), myeloid (CMPs), and granulocyte-macrophage (GMPs) progenitors were found in equal amounts in WT and KO animals. However, we detected a slight but significant decrease in Lin^-^/Kit^+^ (LK) cells, matching the observed decrease in Kit^+^ cells in FL, although total bone marrow cellularity was not significantly decreased (Figure 4E). We also observed a decrease in megakaryocyte-erythroid bi-potent progenitors (MEPs), consistent with the opening and putative activity of the -81kb enh at this stage (Figure 4C-D, Fig.2, and Supplementary Figure 1). Surprisingly, we detected an increase in the percentage of Ter119^+^ population representing differentiating erythroid populations (Figure 4E). We next focused specifically on the Ter119^+^ erythroid compartment and analyzed terminal erythropoiesis as the ratio of immature/mature erythroblasts. Erythroid populations gradually acquire differentiation marker Ter119 and lose CD71 expression as they progress throughout terminal differentiation. The percentage of CD71^high^Ter119^med^ pro-erythroblasts (ProE), Ter119^+^CD71^+^FSC^high^ (EryA), Ter119^+^CD71^+^FSC^low^ cells (late basophilic and poly-chromatic erythroblasts, EryB cells), and Ter-119^+^CD71^−^FSC^low^ cells (orthochromatic erythroblasts and reticulocytes, EryC cells) were within the range of control mice (Figure 4F-G). Accordingly, the erythroid maturation index was similar to controls (Figure 4H). Taken altogether these data indicate that deletion of the -81kb enh does not globally impair hematopoiesis, but affects the percentage of MEPs and of Ter119^+^ erythroid cells without altering the ratio of immature/mature erythroblasts in the adult bone marrow.

**Figure 4:**
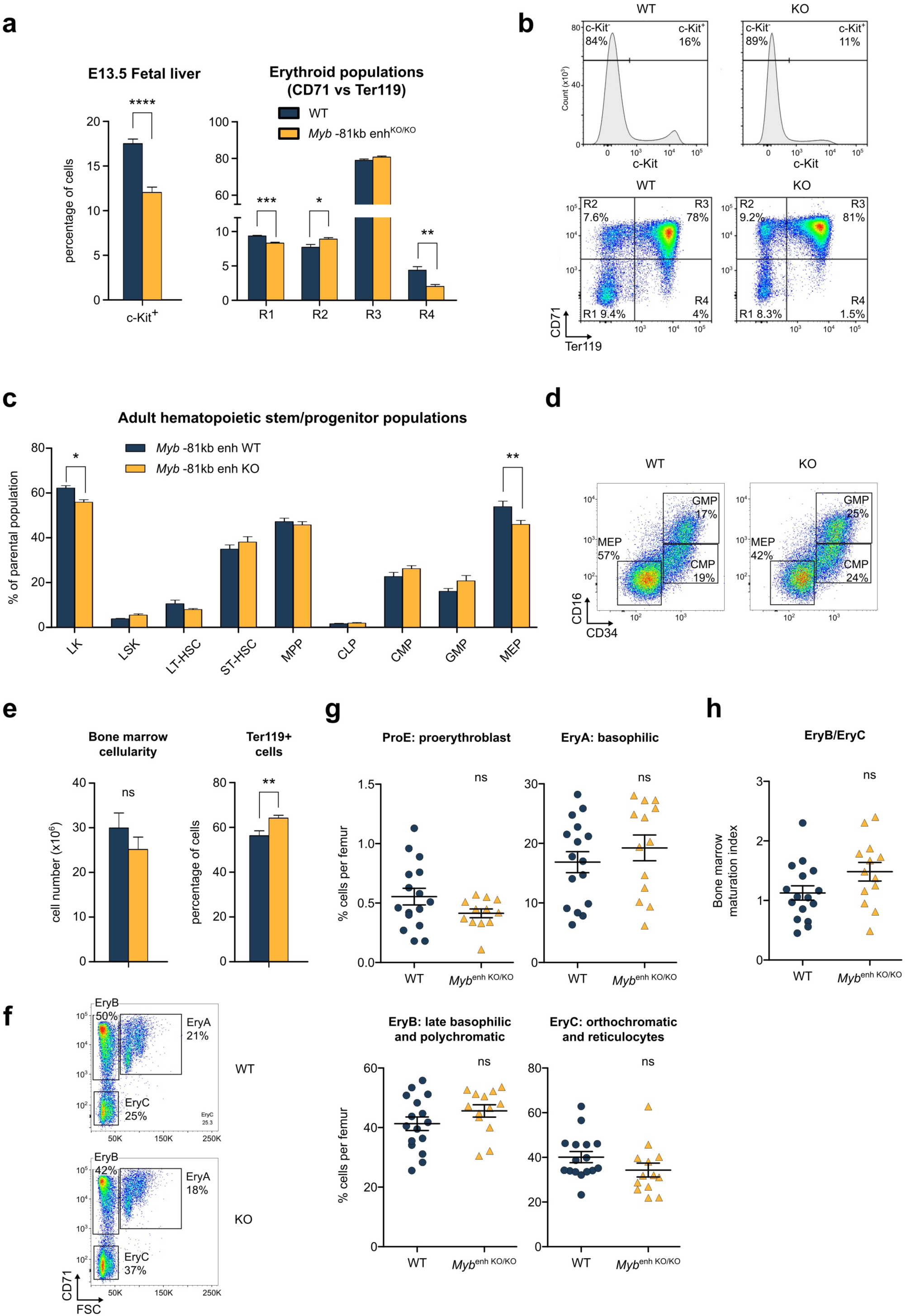
Analysis of fetal liver and bone marrow hematopoiesis of *Myb* -81kb enh^KO^ mice. (**a**) Analysis of c-Kit+ populations and FACS analysis of CD71/Ter119 populations in E13.5 fetal livers from WT and KO mice. (**b**) Representative FACS plots of WT and KO animals. (**c**) FACS analysis of various hematopoietic compartments from WT and KO animals. LK: Lin^-^Kit^+^, LSK: Lin^-^Sca1^+^Kit^+^, LT-HSC: long-term HSCs, ST-HSC: short-term HSCs, MPP: multipotent progenitors, CLP: common lymphoid progenitors, CMP: common myeloid progenitors, GMP: granulocyte/macrophage progenitors, MEP: megakaryocyte/erythroid progenitors. (**d**) representative FACS plots of WT and KO GMP, CMP and MEP populations based on CD34/CD16 surface markers. (**e**) Bone marrow cellularity and frequency of Ter119^+^ erythroid cells. (**f-g**) Erythroblast differentiation in bone marrow evaluated by CD71 and Ter119 staining, with the (**h**) bone marrow erythroid maturation assessed by CD71 and forward scatter area on Ter119+erythroblasts. All data are expressed as the mean ± s.e.m. \**P* < 0.05, ** *P* < 0.01, *** *P* < 0.001, **** *P* < 0.0001.

### Erythroid parameters of *Myb* -81kb enh^KO^ animals and recovery from hemolytic anemia

Since genetic variants at the human -84kb enh are associated with variations in clinical erythroid parameters^56^, we tested whether it is also the case in the mouse. We therefore monitored the erythroid parameters of -81kb enh^KO^ animals from circulating blood sampling in adult (7-10 weeks old) animals. As expected, white blood cell (WBC) counts were unaffected (Supplementary Figure 3). Concerning erythroid cells, whereas red blood cell (RBC) counts, hematocrit (HCT), hemoglobin (Hb) concentration, red cells distribution width (RDW), mean corpuscular Hb (MCH), and MCH concentration (MCHC) were within similar ranges as in control animals, mean corpuscular volume (MCV) values in KO animals were significantly increased (Figure 5A). Increased MCV could be the result of decreased MYB expression (see below) and accelerated erythroid differentiation, as suggested in previous studies^42,43,57,58^, which is compatible with our flow cytometry data showing modified immature/mature erythroblasts ratios (Figure 4B-C). Interestingly, this mimics the human situation where *HBS1L-MYB* intergenic variants associate with increased MCV^42,43,59–63^, indicating that the -81kb enh^KO^ model recapitulates features observed in humans.

**Figure 5:**
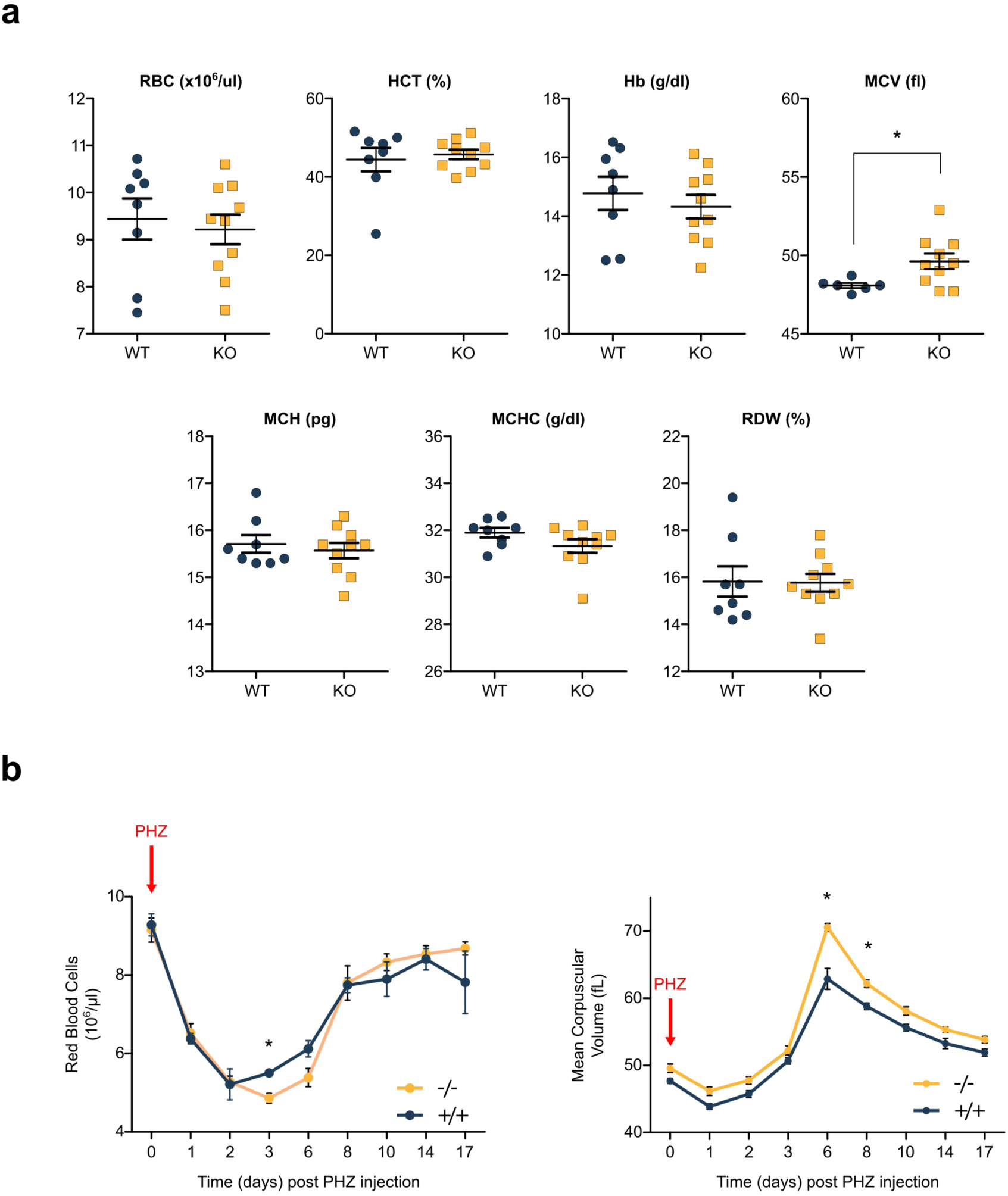
Circulating blood parameters and recovery from hemolytic anemia. (**a**) Erythroid parameters of WT and KO animals measured from circulating blood. RBC: red blood cells, HCT: hematocrit, Hb: hemoglobin, MCV: mean corpuscular volume, MCH: mean corpuscular hematocrit. (**b**) Mice were challenged with phenylhydrazine injections and monitored for recovery over a period of 17 days. Reb blood cell counts and MCV values are shown. All data are expressed as the mean ± s.e.m. \**P* < 0.05.

Since MYB is involved in sustaining erythroid progenitors proliferation, we tested whether -81kb enh deletion alters progenitors capacity to support strong erythroid progenitors expansion following hemolytic challenge. Wild type and -81kb enh^KO^ animals were injected with phenylhydrazine (PHZ) to induce hemolytic anemia and were monitored during the recovery phase. In control mice, a ∼50% reduction in RBC concentration was observed within the first 2 days following the PHZ challenge, followed by a gradual increase starting at day 3 to reach back normal values by day 10 to 14 (Figure 5B). In contrast, KO animals displayed a prolonged period of anemia, with a delayed recovery phase starting only at day 6 post challenge (Figure 5B). However, KO animals managed to fully recover by day 10-14 as in controls. Interestingly, consistent with the blood parameters analysis (Figure 5A), MCV values in KO animals were continuously higher than in controls, both at day 0 and during the whole recovery phase. Taken altogether, these data suggest that the lack of the -81kb distal *Myb* enh impairs progenitor expansion following an acute anemic stress, but does not prevent full recovery. This is suggestive of a reduced proliferative capacity upon *Myb* -81kb enh deletion.

### Decreased *Myb* expression in -81kb enh^KO^ fetal liver cells associates with impaired globin switching and decreased adult globins expression

Since complete *Myb* inactivation leads to an early block of definitive erythroid development at the fetal liver (FL) stage^64^, we monitored *Myb* expression levels in -81kb enh^KO^ sorted FL erythroid populations (Figure 6A). As shown in Figure 6B, *Myb* levels were decreased in immature erythroid progenitors, indicating that the -81kb enh is required to reach maximal expression levels in early erythroid differentiation. In physiological conditions, *Myb* expression dramatically decreases as cells terminally differentiate^41,53,65,66^. We observed a significantly stronger *Myb* silencing in KO animals, supporting a role of the -81kb enh in the fine-tuning of *Myb* regulation in differentiating erythroid cells.

**Figure 6:**
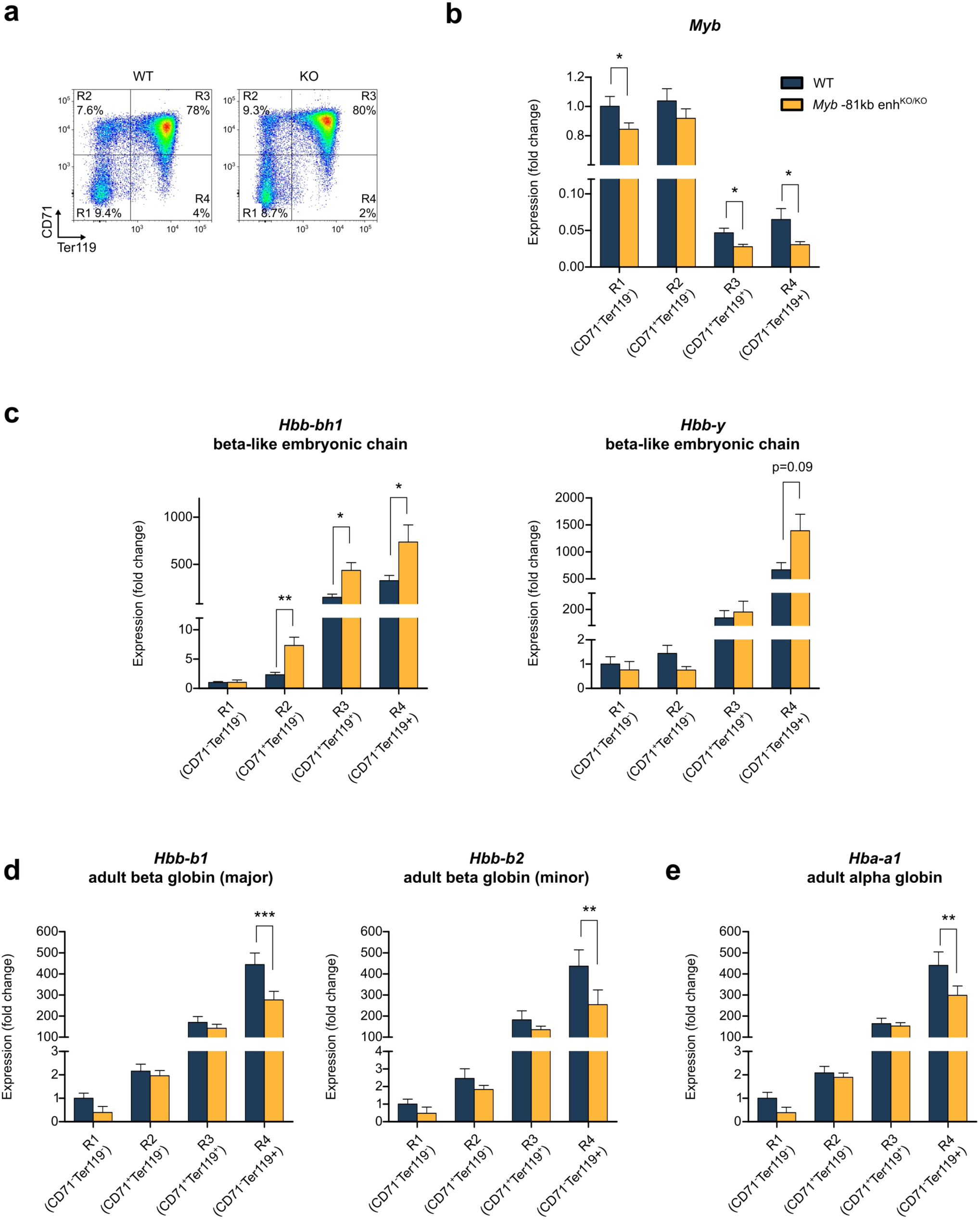
Adult and fetal globin genes expression in *Myb* -81kb enh^KO^ animals. (**a**) E13.5 fetal liver cells were sorted into four populations based on the CD71 and Ter119 surface markers. Sorted cells were used to perform qRT-PCR analyses of (**b**) *Myb*, (**c**) embryonic globins *Hbb-bh1*, *Hbb-y*, (**D**) adult β-globin *Hbb-b1*, *Hbb-b2*, and (**E**) adult α-globin *Hba-a1* genes expression. All data are expressed as the mean ± s.e.m. \**P* < 0.05, \*\**P* < 0.01, \*\*\**P* < 0.001.

Since MYB levels strongly correlate with fetal globin reactivation in humans^42,43,67^, we next analyzed the embryonic beta-like globin genes (*Hbb-bh1*, *Hbb-y*) expression in KO animals. We detected significant upregulation of the hemoglobin Z embryonic beta-like chain *Hbb-bh1* in -81kb enh^KO^ definitive erythroid cells, and a similar trend towards upregulation (although not statistically significant) of the hemoglobin Y embryonic beta-like *Hbb-y* (Figure 6C). This result is in accordance with the increased HBF levels in individuals bearing *HBS1L-MYB* intergenic variants, and indicates that the -81kb enh^KO^ model recapitulates the partial globin switch reversal (or impaired fetal globin repression) seen in humans. Very interestingly, when measuring adult globin genes expression, we observed a consistent and significant decrease in β-globin (*Hbb-b1*, *Hbb-b2*), and α-globin (*Hba-a1*) genes (Figure 6D). This is particularly interesting since excessive amounts of unpaired α-globin proteins in β-thalassemia are a cause of ineffective erythropoiesis, through the formation of toxic aggregates in erythroid cells^68^. Accordingly, reduction of α-globin content and increased fetal β-like globin genes in β-thalassemia are effective in reducing globin chain imbalance and ameliorate disease course. Our result therefore prompted us to investigate whether deletion of the -81kb enh could, at least partially, rescue a thalassemic phenotype in the mouse.

### *Myb* -81kb enh^KO^ ameliorates erythroid parameters *in vivo* in a mouse model of β-thalassemia

To study the role of the *Myb* -81kb enh in hemoglobinopathies, we used a preclinical model of β-thalassemia intermedia, the *Hbb*^th1/th1^ mouse^44^. The *Hbb*^th1/th1^ β-thalassemic mice recapitulate several features of the human disease including chronic anemia and splenomegaly. The -81kb enh KO mice and *Hbb*^th1/th1^ mice were bred to homozygosity and checked for disease-specific erythroid parameters in order to determine if inactivation of the *Myb* -81kb enh ameliorates disease course (Figure 7A). We first noticed that the splenomegaly associated with β-thalassemia in *Hbb*^th1/th1^ mice was significantly reduced in *Hbb*^th1/th1^::*Myb*-81kb enh^KO/KO^ animals (referred to as double KO) (Figure 7B), indicating that -81kb enh KO limits extramedullary erythropoiesis in the *Hbb*^th1/th1^ thalassemic context. Interestingly, we could detect in young adult double KO animals (7-10 weeks old) a slight but significant increase in RBC counts (7,512 ± 0,1446 N=39 in *Hbb*^th1/th1^ animals versus 8,016 ± 0,1068 N=60 in double KO animals) (Figure 7C). Hematocrit levels were also weakly but significantly raised in double KO animals, together with total hemoglobin levels, and mean corpuscular hemoglobin (Figure 7C). Surprisingly, none of these parameters differed significantly from control WT animals in *Myb* -81kb enh KO mice (with the notable exception of the mean corpuscular volume, see also Figure 5), indicating that inactivation of the -81kb enh is phenotypically silent in steady state conditions, but provides a phenotypic advantage in stressed conditions such as β-thalassemia. However, it should be noted that none of the erythroid parameters could be rescued to the levels seen in control animals, indicating that *Myb* -81enh KO on its own is not sufficient to fully rescue disease phenotype in this model of β-thalassemia.

**Figure 7:**
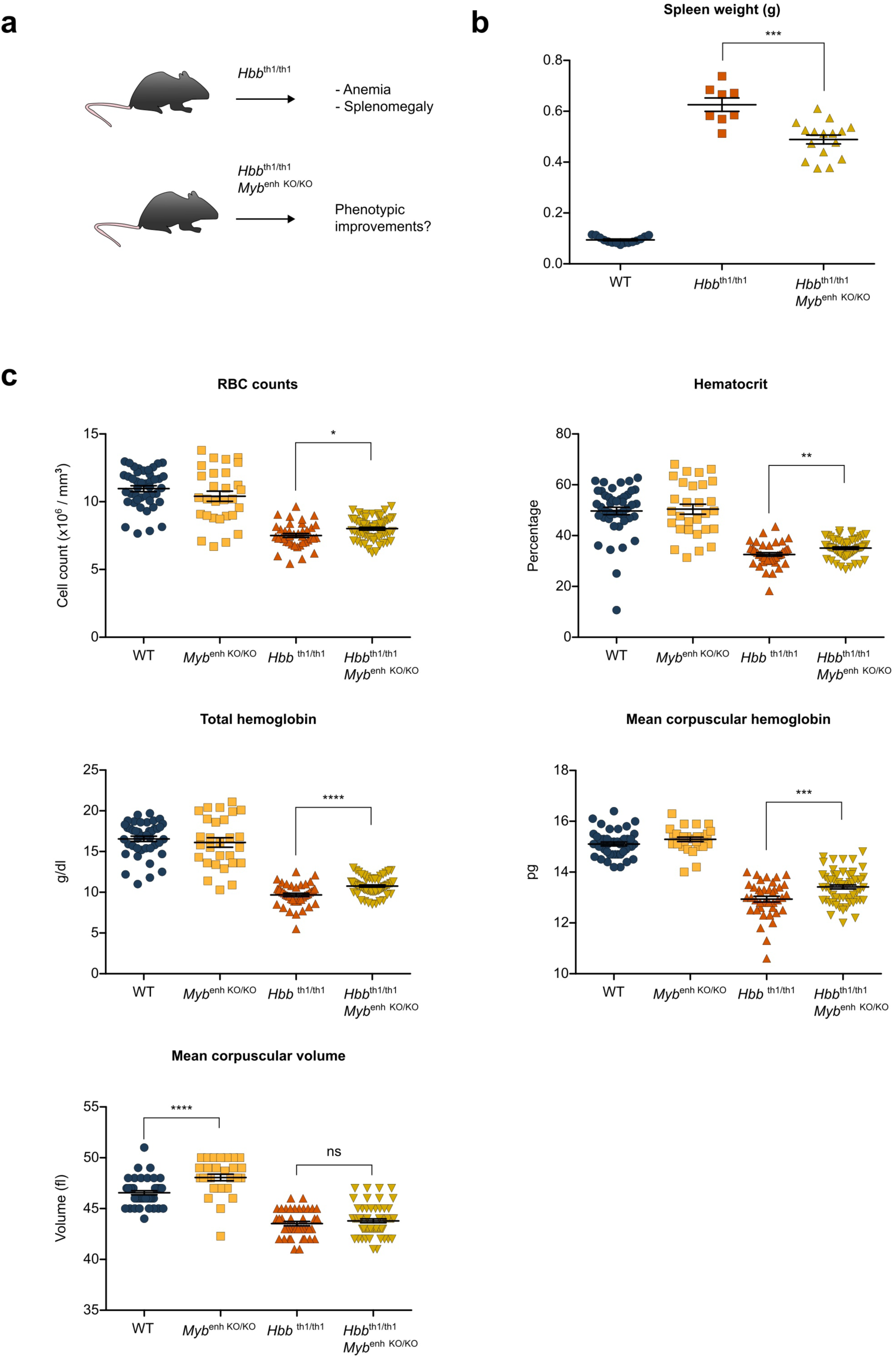
phenotypic impact of *Myb*-81kb enh^KO^ in *Hbb*^th1/th1^ thalassemic mice. (**a**) experimental set-up, (**b**) spleen weight of control, ***Hbb*^th1/th1^, and *Hbb*^th1/th1^::***Myb*-81kb enh^KO^ mice showing decreased splenomegaly. (**c**) Circulating erythroid blood parameters of 7-10 weeks old adult animals from the different genotypes. All data are expressed as the mean ± s.e.m. \**P* < 0.05, ** *P* < 0.01, *** *P* < 0.001, **** *P* < 0.0001.

## Discussion

GWAS studies have been invaluable to identify key genomic loci responsible for disease susceptibility and heritability. However, these studies usually highlight rather large genomic areas and fail to precisely delineate the regulatory regions involved. Genomic variations at the *HBS1L-MYB* locus are such an example. Multiple SNPs in linkage disequilibrium are located in this intergenic region, and are associated with HPFH, a feature ameliorating disease symptom in some of the most common human genetic disorders worldwide, namely sickle cell disease and β-thalassemia. Because common and rare associated variants cover a large region encompassing multiple putative enhancers, functional assessment of the different variants and precise determination of the regulatory elements involved remained enigmatic.

Published data from us and others have shown that the *Myb* -81kb enh, which is the orthologue of the human *MYB* -84kb enh, represents a peculiar enhancer within the locus. First, recent genetic data investigating variants linked to HPFH have robustly identified the human *MYB* -84kb enh as key regulatory site linked to *MYB* regulation and fetal globin reactivation^39,40^. Second, epigenomic data highlighted the mouse -81kb enh/human -84kb enh as being specifically bound by both the GATA1/LDB1 complex and KLF1, a master regulator of terminal erythroid differentiation and globin genes regulation^41,69,70^. Third, saturating mutagenesis coupled to functional analyses highlighted clear *MYB* regulatory potential at the -84kb enh mediated, at least in part, by GATA1 in human cells^19^. Fourth, interestingly, a previous report^71^ and a recent study^45^ reported that the mouse *Myb* -81kb enh harbors the transcriptional initiation site of the *Myrlin* long non-coding RNA, shown to be involved in the cis-regulation of *Myb* expression by recruiting or stabilizing the MLL/COMPASS histone methyltransferase complex and stimulating RNA polymerase pause release to promote *Myb* transcription^45^. Collectively, these observations point at the -81kb/-84kb enh as being a key candidate for the observed erythroid phenotypic variations linked to the top scoring *HBS1L-MYB* variants. We therefore unequivocally tested its function by deleting the -81kb enh sequence from the mouse genome and assessing its functional impact *in vivo* on erythropoiesis, globin genes expression and thalassemia.

We show here that the -81/84kb enh displays erythroid specificity as judged by its epigenetic accessibility profile throughout human and mouse hematopoiesis. Interestingly, its deletion is well tolerated *in vivo* and *Myb* -81kb enh KO animals were viable and fertile contrarily to *Myb* KO mice. This shows that this enhancer is not dominant within the *Myb* locus *in vivo*, and suggests that functional redundancy likely exists between the various *Myb* distal enhancers, buffering the effects of -81kb enh deletion. Alternatively, this may indicate that the -81kb enh is a weak enhancer and has little overall impact on *Myb* expression. Interestingly, in a recent study, in-depth dissection the α-globin enhancer cluster has shown that enhancers with seemingly weak enhancer potential (such as the R4 α-globin enhancer) may actually play key roles as ‘facilitators’ by potentiating the activity of the strong typical enhancers (i.e. R2 enhancer) within the locus, although deletion of the R4 enhancer leads to very little impact on α-globin expression^11,12^. Interestingly, a facilitator-like element was also described in the β-globin enhancer cluster (i.e. HS1), suggesting that such atypical enhancers with weak apparent transcriptional activation capacity may be widespread in mammalian genomes^12^. Accordingly, we found only moderately decreased *Myb* transcript levels in differentiating erythroid cells in KO animals. Whether the -81kb enh belongs to this new class of regulatory elements and potentially act as a facilitator element rather than a typical enhancer remains to be explored. Interestingly, this weak transcriptional effect was also observed in individuals bearing the rs66650371 variant, a 3bp deletion altering LDB1/GATA1 complex binding at the -84kb enh, which in combination with other SNPs in LD show only moderate decrease in *MYB* expression levels^42^. This observation contrasts with previous reports suggesting that Myb -81/-84kb enh is critical for erythroid cell fitness and *Myb* expression^19,45^. A major difference accounting for these discrepancies is the use of cell lines which may be overtly addicted to MYB levels, as reported for transformed leukemic cell lines^72^, or the use of *ex vivo* cultured primary cells that may expand under stressed conditions in the absence of proper tissular supporting microenvironment. By contrast *Myb* -81kb enh KO mice do not show signs of anemia or overt alteration of the erythroid compartment suggesting that the microenvironment may compensate for a slight decrease in erythroid progenitors’ fitness *in vivo*.

It remains unclear why and how variations at this enhancer associate with fetal globin reactivation, but it is interesting to note that as a transcription factor, MYB was shown to target both the globin gene loci themselves, and some of the most potent fetal globin regulators such as BCL11A, KLF1, SOX6^42,66^. A very interesting aspect of our findings is the discovery that in addition to embryonic/fetal globin genes reactivation, *Myb* -81kb enh KO associates with reduction of adult α-globin expression. This is particularly relevant in the case of β-thalassemia as it was shown that excess unpaired α-globin proteins form cytoplasmic aggregates that trap the HSP70 chaperone, which becomes limiting in thalassemic cells. This, in turn, results in the failure to protect the master regulator GATA1 against caspase-mediated cleavage, leading to decreased GATA1 levels and ineffective erythropoiesis^68,73^. Thus, one of the beneficial effects of *Myb* -81kb enh KO may be to partially rescue the α/β globin chains imbalance, thereby ameliorating terminal erythropoiesis. When *Myb* -81kb enh KO animals were crossed with *Hbb*^th1/th1^ thalassemic mice, a marked reduction of splenomegaly was observed, which was accompanied by a significant amelioration of the erythroid parameters, red blood cells counts and hematocrit in young adults, albeit at levels that did not reach wild-type control levels. How the slight reduction of *Myb* expression in -81kb enh KO mice leads to these moderate phenotypic improvements *in vivo* remains to be uncovered. However, MYB is known to be a potent regulator of cell cycle^53,65,74,75^ and under reduced MYB levels, erythroid progenitors display cell cycle kinetic perturbations^19,67,76^, which impact terminal erythropoiesis. Altering cell cycle at these terminal differentiation stages may likely lead to increased fetal hemoglobin as a result of failure to fully switch to adult hemoglobin expression, as proposed in earlier models of HbF reactivation^77^. Accordingly, altered cell cycle progression with accelerated erythroid differentiation were reported in HPFH individuals with decreased MYB expression^42,57,67^. On the other hand, our recent CUT&RUN experiments showed strong enrichment of cell cycle genes and key cell-cycle regulators as direct MYB targets in murine erythroid progenitor cell lines^66^. These studies, together with results reported from hypomorphic *Myb* alleles in the mouse^78,79^, provide evidence for the tight links existing between fine-tuned MYB levels, cell cycle regulation, coordination of globin genes expression, and erythroid differentiation. In our mouse model, reduction of MYB levels likely results in accelerated differentiation due to reduced number of proliferation cycles. This would be consistent with higher MCV values as the erythrocytes are younger red cells, explaining why we consistently observed circulating red blood cells with increased MCV indices in *Myb* -81kb enh KO animals (Figures 5 and 7). Importantly, this reflects what is happening in patients with MYB-associated HPFH^42,43,59–63^.

Whereas the precise perturbations of gene regulatory networks downstream of MYB, together with the genome-wide molecular mechanisms at play remain to be determined, and are outside the scope of this study, our data document for the first time the *in vivo* role of a key *Myb* enhancer in the context of erythroid development, steady state and stressed erythropoiesis, and in disease. It provides a unique model to further dissect the *in vivo* roles of this conserved GWAS-highlighted *Myb* enhancer and to gain deeper mechanistic insights into the role of MYB in the control of terminal erythroid differentiation and β-hemoglobinopathies.

## Methods

### Animals

All experiments were conducted by authorized personnel in accordance with the guidelines of the European Community and approved by the French Ministry of research under approbation numbers APAFIS 18144 v2. The authors complied with the ARRIVE guidelines. *Myb* -81kb enh^KO^ and *Hbb*^th1/th1^ mice were maintained under pathogen-free conditions in our animal facility (RAM-PCEA or RAM-ZEFI). The *Hbb*^th1/th1^ mouse model originated from a natural occurring deletion of the gene encoding β-globin^44^. Mice genotypes were determined by PCR. All mice were on a C57BL/6 background, and both sexes were used without differences between males and females being observed.

### Generation of the *Myb* -81kb enh^KO^ mouse model

Homologous recombination of mouse embryonic stem cells (ESCs) was performed using the targeting construct shown in Figure 3C. Proper targeting was confirmed by PCR and southern blotting. Two independent targeted ESC lines were injected into blastocysts, and one gave germline transmission. F1 mice were bred with mice ubiquitously expressing the Flp recombinase to remove the neomycin resistance marker. These animals were further crossed with mice ubiquitously expressing the Cre recombinase to induce deletion of the *Myb* -81kb enh sequence. Removal of the -81kb enh sequence was confirmed by PCR.

### Blood parameters analysis

Blood samples (50 μL) were collected from facial veins to EDTA-coated tubes. Complete blood counts were measured on scil Vet abc Plus+ (Horiba) according to the manufacturer’s instructions.

### Induction of hemolytic anemia and blood parameters analysis

Mice received sterile phenylhydrazin (PHZ) solution intraperitoneally on day -1 and day 0 with an 80 mg/kg body weight dose. Blood samples (50 μL) were collected from day 0 to day 17, from facial veins to heparin-coated tubes to monitor the recovery of erythropoiesis post PHZ injection.

### Flow cytometry

Flow cytometry analyses were performed on bone marrow cells (from tibias and femurs) or spleen cells from adult mice and fetal liver cells from E13.5 embryos. Antibodies used were listed in Table 1. Cells were analyzed with a BD FACS-Quanto II or sorted with a BD FACS Aria.

**Table 1:** Flow cytometry antibodies used.

| Antidoby | Reference | Supplier |
| --- | --- | --- |
| CD16/CD32-PerCP-Cy5.5 | 45-0161-82 | eBioscience™/ Invitrogen |
| CD34-FITC | 11-0341-82 | eBioscience™/ Invitrogen |
| CD48-PE-Cy7 | 103424 | Biolegend |
| CD71-PE | 553267 | BD Biosciences |
| CD117(c-Kit)[2B8]-APC | 17-1171-83 | eBioscience™/ Invitrogen |
| CD117(c-Kit)[ACK2]-APC | 17-1172-82 | eBioscience™/ Invitrogen |
| CD117(c-Kit)[2B8]-APC-eF780 | 47-1171-82 | eBioscience™/ Invitrogen |
| CD135(Flt3)-PE | 12-1351-81 | eBioscience™/ Invitrogen |
| CD150-APC | 115910 | Biolegend |
| FVD-eF506 | 65-0866-18 | eBioscience™/ Invitrogen |
| Sca1-APC | 17-5981-82 | eBioscience™/ Invitrogen |
| Sca1-PerCP-Cy5.5 | 45-5981-82 | eBioscience™/ Invitrogen |
| Streptavidin-eF450 | 48-4317-82 | eBioscience™/ Invitrogen |
| Ter119-eFluor450 | 48-5921-82 | eBiosciences / Invitrogen |

### RNA preparation and real-time polymerase chain reaction analysis

Total RNA from sorted cells were extracted using HP RNA isolation kit (Roche) according to manufacturer’s instructions. RNA were primed with oligo(dT) and reverse transcribed with SuperScript II (Invitrogen). Real-time polymerase chain reaction was performed using LightCycler® 480 SYBR Green I Master mix (Roche). The sequences of the primers used for PCR amplification are listed in Table 2.

**Table 2:**
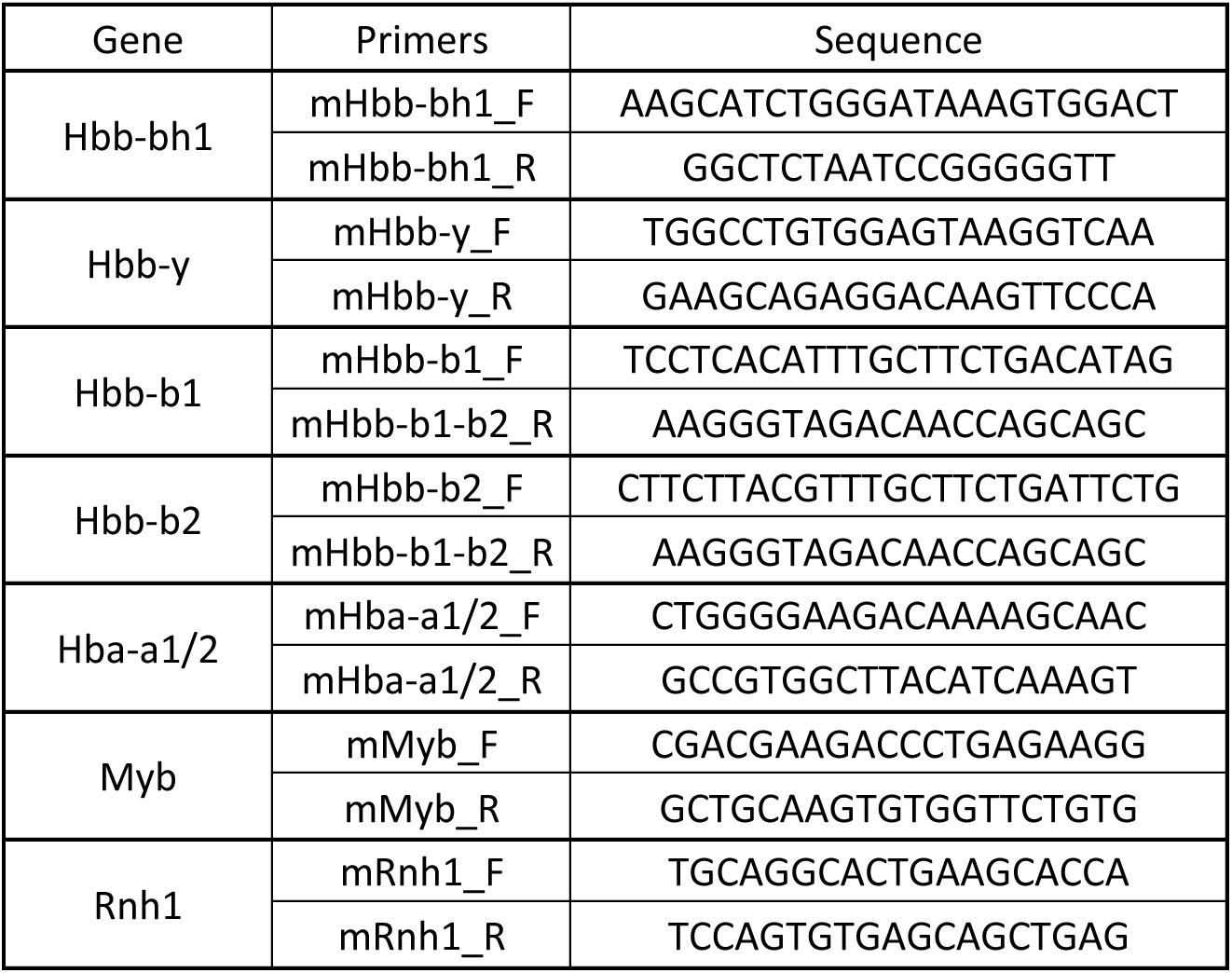
qRT-PCR primers used.

### Statistical analyses

Statistical analysis was performed using GraphPad Prism 5.0.0 software. Unpaired Student’s t-test was used to determine the statistical significance of the difference between two groups. Results with p-values ≤ 0.05 were considered as significant (* = *P* < 0.05, ** = *P* < 0.01, ***= *P* < 0.001, **** = *P* < 0.0001, ns = not significant).

## Acknowledgements

This work was supported by the France 2030 program through the Idex Université Paris Cité (ANR-18-IDEX-0001), the Laboratory of Excellence GR-Ex [ANR-11-LABX-0051; “Investissements d’avenir” program of the French National Research Agency, reference ANR-11-IDEX-0005-02]; the Labex EpiGenMed [“Investissements d’avenir” program, reference ANR-10-LABX-12-01]; the Fondation pour la Recherche Médicale (Equipe FRM DEQ20180339221), and the SIRIC Montpellier Cancer, Grant « INCa-DGOS-INSERM-ITMO Cancer_ 18004 ». We are grateful to the animal facility platform RAM-PCEA and IGMM animal experimentation platform RAM-ZEFI which are part of the “Réseau des Animaleries Montpelliéraines” RAMIBiSA Facility for animal experiments. We acknowledge the “Réseau d’Histologie Expérimentale de Montpellier” -RHEM facility (supported by SIRIC Montpellier Cancer Grant INCa_Inserm_DGOS_12553, the european regional development foundation and the occitanian region (FEDER-FSE 2014-2020 Languedoc Roussillon), REACT-EU (Recovery Assistance for Cohesion and the Territories of Europe), IBiSA and Ligue contre le cancer) for processing animal tissues, histology technics and expertise. We acknowledge the imaging facility MRI, member of the national infrastructure France-BioImaging infrastructure supported by the French National Research Agency (ANR-10-INBS-04, «Investments for the future»). We wish to thank Stéphanie Viala and Myriam Boyer from MRI platform for the help with flow cytometry and cell sorting.

## Author contributions

Conceptualization, E.S.; Methodology, V.D., C.A.S., T.S., R.J., A.M., M.L.A., F.G. and E.S.; Investigation, V.D., T.S., C.O., V.B., R.J., A.M., M.L.A., M.S., C.A.S., F.G.; Data Curation, V.D., M.S., T.S., E.S.; Writing – Original Draft, E.S.; Funding Acquisition, E.S.

## Data availability statement

Data and material of this study can be made available upon reasonable request to

## Competing interests

The authors declare no competing interests.

## Notes

### Competing Interest Statement

The authors have declared no competing interest.

## References

1. Maurano, M. T. et al. Systematic Localization of Common Disease-Associated Variation in Regulatory DNA. Science 337, 1190–1195 (2012).

2. Vierstra, J. et al. Global reference mapping of human transcription factor footprints. Nature 583, 729–736 (2020).

3. Long, H. K., Prescott, S. L. & Wysocka, J. Ever-Changing Landscapes: Transcriptional Enhancers in Development and Evolution. Cell 167, 1170–1187 (2016).

4. Ray-Jones, H. & Spivakov, M. Transcriptional enhancers and their communication with gene promoters. Cell. Mol. Life Sci. 78, 6453–6485 (2021).

5. Andrieu-Soler, C. & Soler, E. Erythroid Cell Research: 3D Chromatin, Transcription Factors and Beyond. IJMS 23, 6149 (2022).

6. Lim, F. et al. Affinity-optimizing enhancer variants disrupt development. Nature 626, 151– 159 (2024).

7. Robson, M. I., Ringel, A. R. & Mundlos, S. Regulatory Landscaping: How Enhancer-Promoter Communication Is Sculpted in 3D. Molecular Cell 74, 1110–1122 (2019).

8. Bresnick, E. H. & Johnson, K. D. Blood disease–causing and –suppressing transcriptional enhancers: general principles and GATA2 mechanisms. Blood Advances 3, 2045–2056 (2019).

9. Grosveld, F., van Assendelft, G. B., Greaves, D. R. & Kollias, G. Position-independent, high-level expression of the human beta-globin gene in transgenic mice. Cell 51, 975–985 (1987).

10. Grosveld, F., Van Staalduinen, J. & Stadhouders, R. Transcriptional Regulation by (Super)Enhancers: From Discovery to Mechanisms. Annu. Rev. Genom. Hum. Genet. 22, 127–146 (2021).

11. Hay, D. et al. Genetic dissection of the α-globin super-enhancer in vivo. Nat Genet 48, 895–903 (2016).

12. Blayney, J. W. et al. Super-enhancers include classical enhancers and facilitators to fully activate gene expression. Cell 186, 5826–5839.e18 (2023).

13. Liu, X. et al. In Situ Capture of Chromatin Interactions by Biotinylated dCas9. Cell 170, 1028–1043.e19 (2017).

14. Deng, W. et al. Controlling Long-Range Genomic Interactions at a Native Locus by Targeted Tethering of a Looping Factor. Cell 149, 1233–1244 (2012).

15. Deng, W. et al. Reactivation of Developmentally Silenced Globin Genes by Forced Chromatin Looping. Cell 158, 849–860 (2014).

16. Krivega, I., Dale, R. K. & Dean, A. Role of LDB1 in the transition from chromatin looping to transcription activation. Genes Dev. 28, 1278–1290 (2014).

17. Bozhilov, Y. K. et al. A gain-of-function single nucleotide variant creates a new promoter which acts as an orientation-dependent enhancer-blocker. Nat Commun 12, 3806 (2021).

18. Sher, F. et al. Rational targeting of a NuRD subcomplex guided by comprehensive in situ mutagenesis. Nat Genet 51, 1149–1159 (2019).

19. Canver, M. C. et al. Variant-aware saturating mutagenesis using multiple Cas9 nucleases identifies regulatory elements at trait-associated loci. Nat Genet 49, 625–634 (2017).

20. Martin-Rufino, J. D. et al. Transcription factor networks disproportionately enrich for heritability of blood cell phenotypes. Preprint at 10.1101/2024.09.09.611392 (2024).

21. Cheng, L. et al. Single-nucleotide-level mapping of DNA regulatory elements that control fetal hemoglobin expression. Nat Genet 53, 869–880 (2021).

22. Fontana, L., Alahouzou, Z., Miccio, A. & Antoniou, P. Epigenetic Regulation of β-Globin Genes and the Potential to Treat Hemoglobinopathies through Epigenome Editing. Genes 14, 577 (2023).

23. Liu, N. et al. Direct Promoter Repression by BCL11A Controls the Fetal to Adult Hemoglobin Switch. Cell 173, 430–442.e17 (2018).

24. Masuda, T. et al. Transcription factors LRF and BCL11A independently repress expression of fetal hemoglobin. Science 351, 285–289 (2016).

25. Qin, K. et al. Dual function NFI factors control fetal hemoglobin silencing in adult erythroid cells. Nat Genet 54, 874–884 (2022).

26. Chaand, M. et al. Erythroid lineage chromatin accessibility maps facilitate identification and validation of NFIX as a fetal hemoglobin repressor. Commun Biol 6, 640 (2023).

27. Soler, E. et al. The genome-wide dynamics of the binding of Ldb1 complexes during erythroid differentiation. Genes Dev. 24, 277–289 (2010).

28. Palstra, R.-J. et al. The β-globin nuclear compartment in development and erythroid differentiation. Nat Genet 35, 190–194 (2003).

29. Bank, A. Regulation of human fetal hemoglobin: new players, new complexities. Blood 107, 435–443 (2006).

30. Thein, S. L. Genetic association studies in β-hemoglobinopathies. Hematology 2013, 354–361 (2013).

31. Andrieu-Soler, C. & Soler, E. When basic science reaches into rational therapeutic design: from historical to novel leads for the treatment of β-globinopathies. Curr Opin Hematol 27, 141–148 (2020).

32. Menzel, S. et al. A QTL influencing F cell production maps to a gene encoding a zinc-finger protein on chromosome 2p15. Nat Genet 39, 1197–1199 (2007).

33. Thein, S. L., et al. Intergenic variants of HBS1L-MYB are responsible for a major quantitative trait locus on chromosome 6q23 influencing fetal hemoglobin levels in adults. Proc. Natl. Acad. Sci. U.S.A. 104, 11346–11351 (2007).

34. Uda, M., et al. Genome-wide association study shows BCL11A associated with persistent fetal hemoglobin and amelioration of the phenotype of β-thalassemia. Proc. Natl. Acad. Sci. U.S.A. 105, 1620–1625 (2008).

35. Lettre, G., et al. DNA polymorphisms at the BCL11A, HBS1L-MYB, and β-globin loci associate with fetal hemoglobin levels and pain crises in sickle cell disease. Proc. Natl. Acad. Sci. U.S.A. 105, 11869–11874 (2008).

36. Galarneau, G. et al. Fine-mapping at three loci known to affect fetal hemoglobin levels explains additional genetic variation. Nat Genet 42, 1049–1051 (2010).

37. Farrell, J. J. et al. A 3-bp deletion in the HBS1L-MYB intergenic region on chromosome 6q23 is associated with HbF expression. Blood 117, 4935–4945 (2011).

38. Danjou, F. et al. Genome-wide association analyses based on whole-genome sequencing in Sardinia provide insights into regulation of hemoglobin levels. Nat Genet 47, 1264–1271 (2015).

39. Ojewunmi, O. O. et al. The genetic dissection of fetal haemoglobin persistence in sickle cell disease in Nigeria. Human Molecular Genetics 33, 919–929 (2024).

40. Cato, L. D. et al. Genetic regulation of fetal hemoglobin across global populations. Preprint at 10.1101/2023.03.24.23287659 (2023).

41. Stadhouders, R. et al. Dynamic long-range chromatin interactions control Myb proto-oncogene transcription during erythroid development. EMBO J 31, 986–999 (2012).

42. Stadhouders, R. et al. HBS1L-MYB intergenic variants modulate fetal hemoglobin via long-range MYB enhancers. J Clin Invest 124, 1699–1710 (2014).

43. Menzel, S. et al. The HBS1L-MYB intergenic region on chromosome 6q23.3 influences erythrocyte, platelet, and monocyte counts in humans. Blood 110, 3624–3626 (2007).

44. Skow, L. C. et al. A mouse model for β-thalassemia. Cell 34, 1043–1052 (1983).

45. Kim, J. et al. An enhancer RNA recruits KMT2A to regulate transcription of Myb. Cell Reports 43, 114378 (2024).

46. Papadopoulos, G. L. et al. GATA-1 genome-wide occupancy associates with distinct epigenetic profiles in mouse fetal liver erythropoiesis. Nucleic Acids Research 41, 4938– 4948 (2013).

47. Schulz, V. P. et al. A Unique Epigenomic Landscape Defines Human Erythropoiesis. Cell Reports 28, 2996–3009.e7 (2019).

48. Ludwig, L. S. et al. Transcriptional States and Chromatin Accessibility Underlying Human Erythropoiesis. Cell Reports 27, 3228–3240.e7 (2019).

49. Lieu, Y. K. & Reddy, E. P. Conditional c-myb knockout in adult hematopoietic stem cells leads to loss of self-renewal due to impaired proliferation and accelerated differentiation. Proc Natl Acad Sci U S A 106, 21689–21694 (2009).

50. Bender, T. P., Kremer, C. S., Kraus, M., Buch, T. & Rajewsky, K. Critical functions for c-Myb at three checkpoints during thymocyte development. Nat Immunol 5, 721–729 (2004).

51. Thomas, M. D., Kremer, C. S., Ravichandran, K. S., Rajewsky, K. & Bender, T. P. c-Myb Is Critical for B Cell Development and Maintenance of Follicular B Cells. Immunity 23, 275– 286 (2005).

52. Wang, X., Angelis, N. & Thein, S. L. MYB – A regulatory factor in hematopoiesis. Gene 665, 6–17 (2018).

53. Ramsay, R. G. & Gonda, T. J. MYB function in normal and cancer cells. Nat Rev Cancer 8, 523–534 (2008).

54. Corces, M. R. et al. Lineage-specific and single-cell chromatin accessibility charts human hematopoiesis and leukemia evolution. Nat Genet 48, 1193–1203 (2016).

55. Xiang, G. et al. An integrative view of the regulatory and transcriptional landscapes in mouse hematopoiesis. Genome Res. 30, 472–484 (2020).

56. Tumburu, L. & Thein, S. L. Genetic control of erythropoiesis. Current Opinion in Hematology 24, 173–182 (2017).

57. Jiang, J. et al. cMYB is involved in the regulation of fetal hemoglobin production in adults. Blood 108, 1077–1083 (2006).

58. Mukai, H. Y. et al. Transgene Insertion in Proximity to the c-*myb* Gene Disrupts Erythroid-Megakaryocytic Lineage Bifurcation. Molecular and Cellular Biology 26, 7953– 7965 (2006).

59. Ganesh, S. K. et al. Multiple loci influence erythrocyte phenotypes in the CHARGE Consortium. Nat Genet 41, 1191–1198 (2009).

60. Soranzo, N. et al. A genome-wide meta-analysis identifies 22 loci associated with eight hematological parameters in the HaemGen consortium. Nat Genet 41, 1182–1190 (2009).

61. Kamatani, Y. et al. Genome-wide association study of hematological and biochemical traits in a Japanese population. Nat Genet 42, 210–215 (2010).

62. Van Der Harst, P. et al. Seventy-five genetic loci influencing the human red blood cell. Nature 492, 369–375 (2012).

63. Kanai, M. et al. Genetic analysis of quantitative traits in the Japanese population links cell types to complex human diseases. Nat Genet 50, 390–400 (2018).

64. Mucenski, M. L. et al. A functional c-myb gene is required for normal murine fetal hepatic hematopoiesis. Cell 65, 677–689 (1991).

65. Oh, I.-H. & Reddy, E. P. The myb gene family in cell growth, differentiation and apoptosis. Oncogene 18, 3017–3033 (1999).

66. Deleuze, V. et al. Efficient genome editing in erythroid cells unveils novel MYB target genes and regulatory functions. iScience 26, 107641 (2023).

67. Sankaran, V. G., et al. MicroRNA-15a and -16-1 act via MYB to elevate fetal hemoglobin expression in human trisomy 13. Proc. Natl. Acad. Sci. U.S.A. 108, 1519–1524 (2011).

68. Arlet, J.-B. et al. HSP70 sequestration by free α-globin promotes ineffective erythropoiesis in β-thalassaemia. Nature 514, 242–246 (2014).

69. Tallack, M. R. et al. A global role for KLF1 in erythropoiesis revealed by ChIP-seq in primary erythroid cells. Genome Res. 20, 1052–1063 (2010).

70. Tallack, M. R. & Perkins, A. C. KLF1 directly coordinates almost all aspects of terminal erythroid differentiation. IUBMB Life 62, 886–890 (2010).

71. Morrison, T. A. et al. A long noncoding RNA from the HBS1L-MYB intergenic region on chr6q23 regulates human fetal hemoglobin expression. *Blood Cells*, Molecules, and Diseases 69, 1–9 (2018).

72. Pacini, C. et al. A comprehensive clinically informed map of dependencies in cancer cells and framework for target prioritization. Cancer Cell 42, 301–316.e9 (2024).

73. Guillem, F. et al. XPO1 regulates erythroid differentiation and is a new target for the treatment of β-thalassemia. haematol 105, 2240–2249 (2020).

74. Nakata, Y. et al. c-Myb Contributes to G_2_ /M Cell Cycle Transition in Human Hematopoietic Cells by Direct Regulation of Cyclin B1 Expression. Molecular and Cellular Biology 27, 2048–2058 (2007).

75. Cicirò, Y. & Sala, A. MYB oncoproteins: emerging players and potential therapeutic targets in human cancer. Oncogenesis 10, 19 (2021).

76. Bianchi, E. et al. c-myb supports erythropoiesis through the transactivation of KLF1 and LMO2 expression. Blood 116, e99–e110 (2010).

77. Stamatoyannopoulos, G., Veith, R., Galanello, R. & Papayannopoulou, Th. Hb F Production in Stressed Erythropoiesis: Observations and Kinetic Modelsa^a^. Annals of the New York Academy of Sciences 445, 188–197 (1985).

78. Emambokus, N. Progression through key stages of haemopoiesis is dependent on distinct threshold levels of c-Myb. The EMBO Journal 22, 4478–4488 (2003).

79. Vegiopoulos, A., García, P., Emambokus, N. & Frampton, J. Coordination of erythropoiesis by the transcription factor c-Myb. Blood 107, 4703–4710 (2006).

80. Huang, S. et al. Mutations in linker-2 of KLF1 impair expression of membrane transporters and cytoskeletal proteins causing hemolysis. Nat Commun 15, 7019 (2024).

81. Downes, D. J. et al. Capture-C: a modular and flexible approach for high-resolution chromosome conformation capture. Nat Protoc 17, 445–475 (2022).

82. Stadhouders, R. et al. Multiplexed chromosome conformation capture sequencing for rapid genome-scale high-resolution detection of long-range chromatin interactions. Nat Protoc 8, 509–524 (2013).

83. Thongjuea, S., Stadhouders, R., Grosveld, F. G., Soler, E. & Lenhard, B. r3Cseq: an R/Bioconductor package for the discovery of long-range genomic interactions from chromosome conformation capture and next-generation sequencing data. Nucleic Acids Res 41, e132 (2013).

84. Brouwer, R. W. W., Van Den Hout, M. C. G. N., Van IJcken, W. F. J., Soler, E. & Stadhouders, R. Unbiased Interrogation of 3D Genome Topology Using Chromosome Conformation Capture Coupled to High-Throughput Sequencing (4C-Seq). in Eukaryotic Transcriptional and Post-Transcriptional Gene Expression Regulation (eds. Wajapeyee, N. & Gupta, R.) vol. 1507 199–220 (Springer New York, New York, NY, 2017).

