## Supplementary figures for "*In vivo* deletion of a GWAS-identified *Myb* distal enhancer acts on *Myb* expression, globin switching, and clinical erythroid parameters in β-thalassemia"

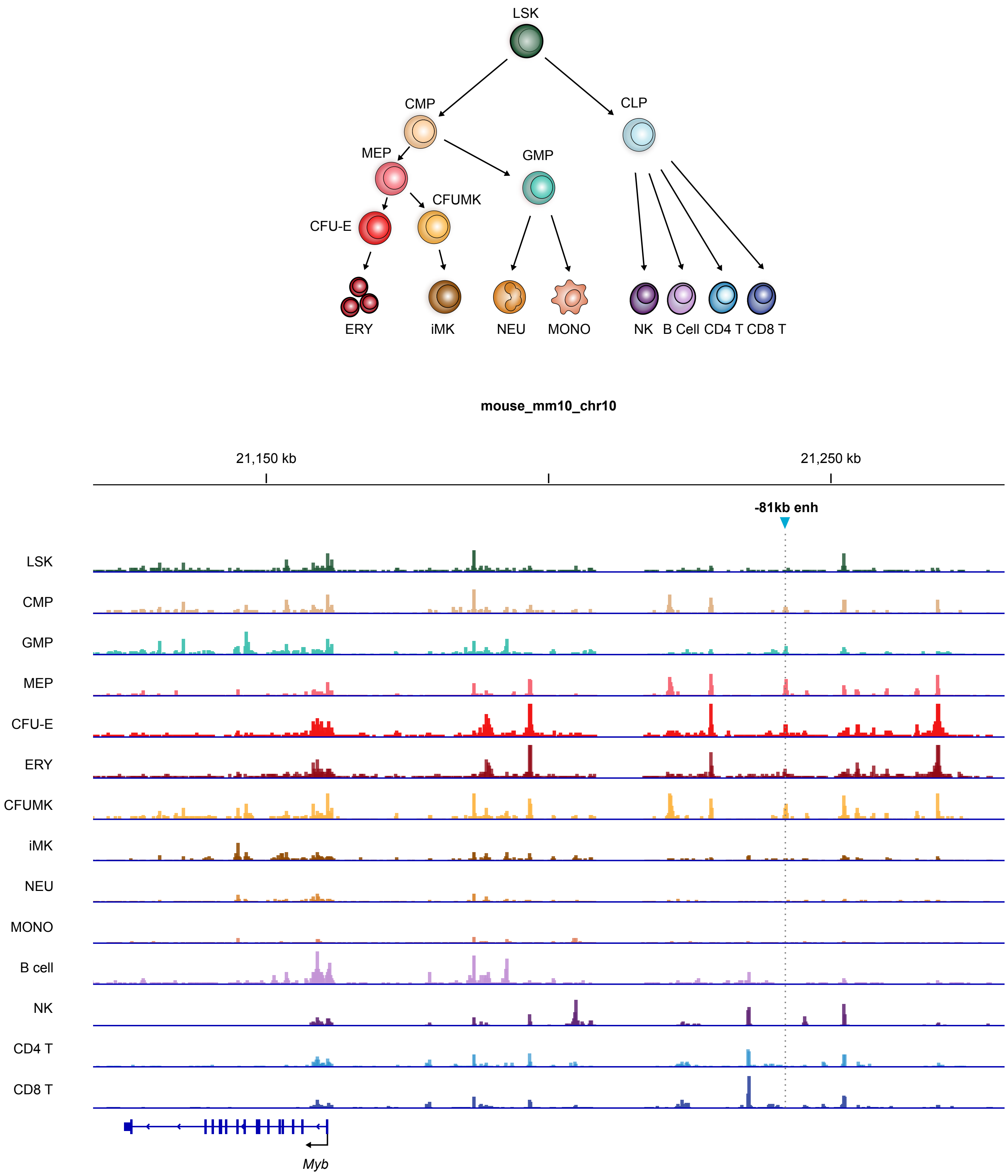


**Supplementary Figure S1 : The Myb chromatin landscape in mouse hematopoietic cells.**

ATAC-Seq signals showing open chromatin regions at the mouse Myb locus from various mouse hematopoietic populations[55] are depicted, showing Myb -81kb enhancer chromatin opening being restricted to erythro-myeloid cells, MEP, erythroid and megakaryocytic progenitors.

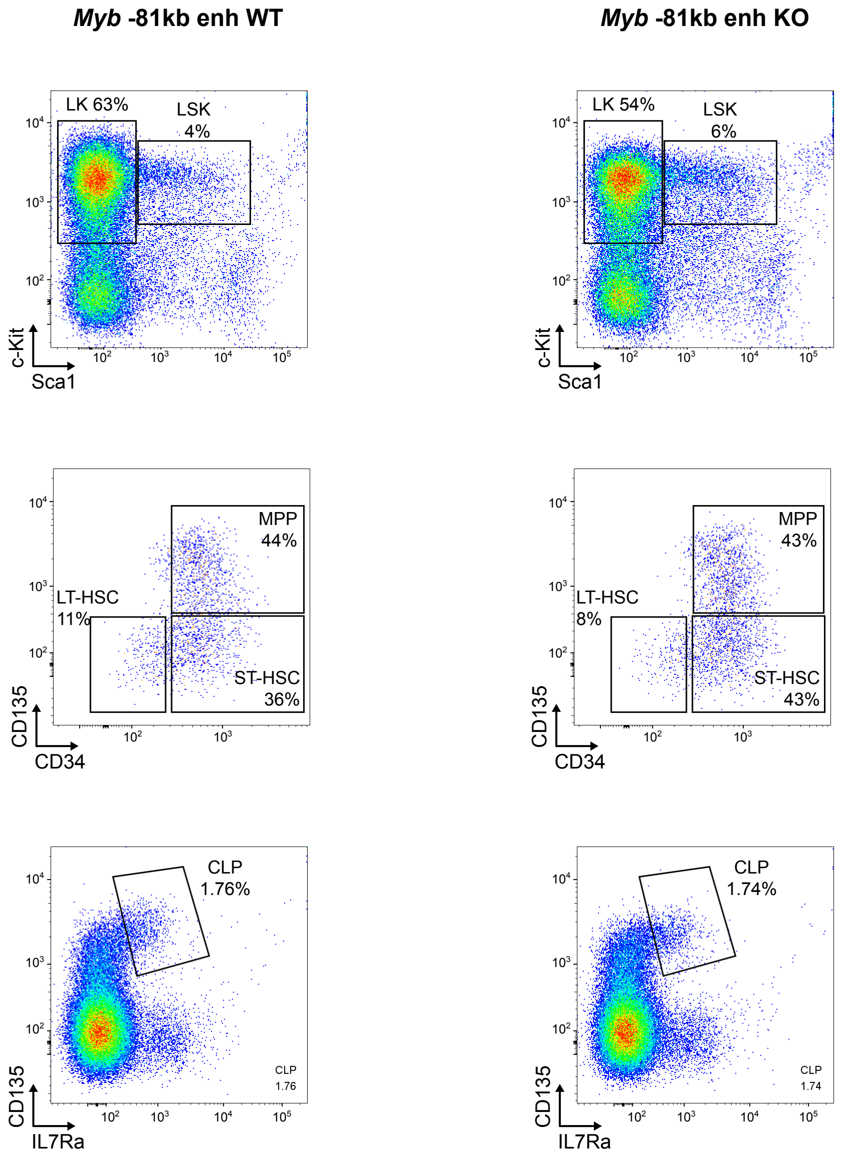


**Supplementary Figure S2 : Detailed analysis of bone marrow hematopoiesis in Myb -81kb enhKO mice.**

Representative FACS plots of various hematopoietic compartments from WT and KO animals. LK: Lin-Kit+, LSK: Lin-Sca1+Kit+, based on Sca1/c-Kit surface markers. LT-HSC: long-term HSCs, ST-HSC: short-term HSCs, MPP: multipotent progenitors, based on CD34/CD135 surface markers. CLP: common lymphoid progenitors, based on IL7Ra/CD135 surface markers.

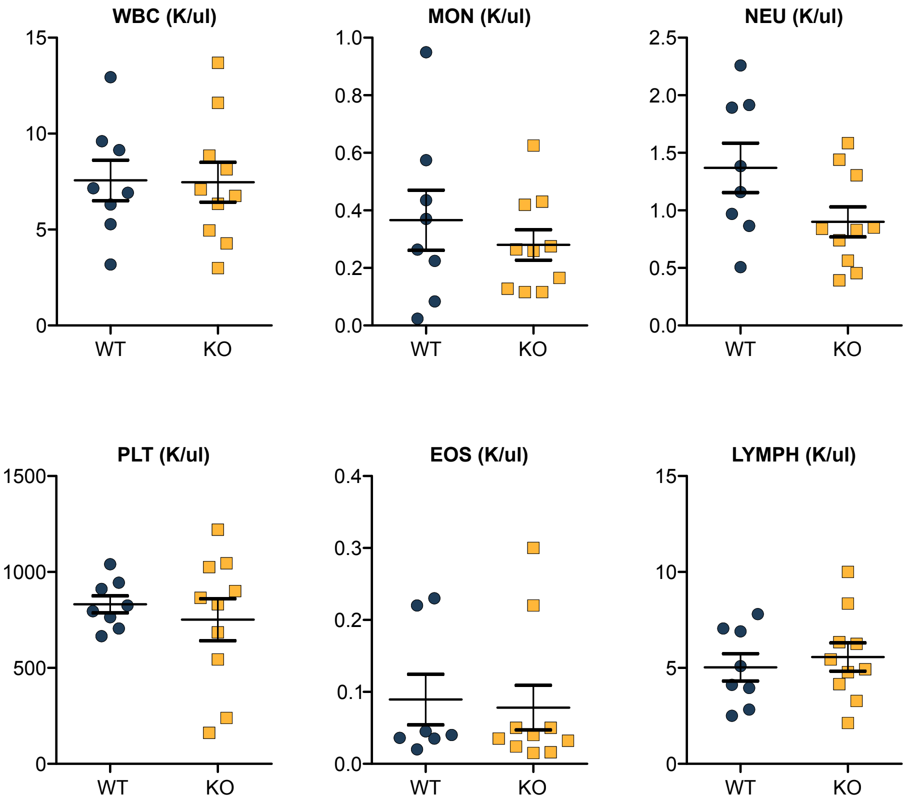


**Supplementary Figure S3 : Complete blood count of Myb -81kb enhKO mice.**

Complete blood cells count of WT and KO animals measured from circulating blood. WBC: white blood cells, MON: monocytes, NEU: neutrophils, PLT: platelets, EOS: eosinophils, LYMPH: lymphocytes.
